# Egalitarian binding partners, Dynein Light Chain and Bicaudal-D, act sequentially to link mRNA to the Dynein motor

**DOI:** 10.1101/550277

**Authors:** Chandler H. Goldman, Rajalakshmi Veeranan-Karmegam, Hannah Neiswender, Graydon B. Gonsalvez

## Abstract

A widely conserved mechanism of polarity establishment is the localization of mRNA to specific cellular regions. While it is clear that many mRNAs are transported to their destinations along microtubule tracks, much less is known regarding the mechanism by which these mRNAs are linked to microtubule motors. The RNA binding protein Egalitarian (Egl) is necessary for localization of several mRNAs in *Drosophila* oocytes and embryos. In addition to binding RNA, Egl also interacts with Dynein light chain (Dlc) and Bicaudal-D (BicD). The role of Dlc and BicD in mRNA localization has remained elusive. Like Egl, both proteins are required for oocyte specification. Null alleles in these genes result in an oogenesis block. In this report, we used an innovative approach to overcome the oogenesis block. Our findings reveal that the primary function of Dlc is to promote Egl dimerization. Loss of dimerization compromises the ability of Egl to bind RNA. Consequently, Egl is not bound to cargo, and is not able to efficiently associate with BicD and the Dynein motor. Our results therefore identify the key molecular steps required for assembling a localization competent mRNP.

## Introduction

Recent studies have shown that mRNA localization is a prevalent, potent, and conserved mechanism used by cells to regulate protein sorting (Lecuyer et al., 2007; Mili et al., 2008; Cajigas et al., 2012; Ryder et al., 2018). Although there are several pathways by which mRNAs are localized, active motor-based transport along cytoskeletal filaments is a commonly used mechanism. In *Drosophila* oocytes and embryos, numerous mRNAs are localized to distinct intracellular sites (Goldman et al., 2017). For many of these mRNAs, localization requires the activity of the minus-end directed microtubule motor, cytoplasmic Dynein (hereafter referred to as Dynein) (Kardon et al., 2009; Suter, 2018). An unanswered question in the field pertains to the mechanism by which Dynein is able to bind these mRNAs in the complex and crowded environment of the cell.

Egalitarian (Egl) is an RNA binding protein that associates with several mRNAs that are localized in a Dynein-dependent manner (Dienstbier et al., 2009). Egl also interacts with Dynein light chain/LC8 (Dlc) and Bicaudal-D (BicD) (Mach et al., 1997; Navarro et al., 2004). Dlc is a component of the Dynein motor and BicD interacts with Dynactin, a critical regulator of Dynein (Hoogenraad et al., 2001; Rapali et al., 2011). Thus, in theory, both of Egl’s binding partners are capable of linking mRNA bound Egl with the Dynein motor. Recent in vitro studies have shown that a minimal complex consisting of RNA, Egl, BicD, Dynactin, and Dynein is sufficient for transport (McClintock et al., 2018; Sladewski et al., 2018). Whether or not this complex is sufficient for mRNA localization in vivo, however, is not known.

Egl, Dlc, BicD, and components of the Dynein motor are required for oocyte specification (Mach et al., 1997; McGrail et al., 1997; Navarro et al., 2004). Null mutants in these genes result in an oogenesis block. As such, mRNA localization, which is primarily studied in mid and late stage egg chambers and early embryos, cannot be analyzed using null alleles. We recently overcame this limitation by using an shRNA depletion strategy (Sanghavi et al., 2016). This approach revealed an essential role for Egl in the localization of *oskar* (*osk*), *bicoid* (*bcd*) and *gurken* (*grk*) mRNAs (Sanghavi et al., 2016). In this study, we used a similar strategy to define the mechanism by which Egl, Dlc, and BicD function to link mRNAs with Dynein. Our results reveal a step-wise assembly pathway for building a localization competent mRNP.

## Results

Egl contains binding sites for BicD and Dlc on opposite ends of the protein and a central region that is essential for RNA binding (Fig. 1A) (Mach et al., 1997; Navarro et al., 2004; Dienstbier et al., 2009). Point mutations have been identified within Egl that disrupt either Dlc or BicD binding. For instance, the *egl2pt* allele contains two mutations that eliminate Dlc interaction (Navarro et al., 2004). The *egl4e* allele contains a single mutation that compromises BicD binding (Mach et al., 1997). However, as with null alleles of *egl*, flies that are homozygous for *egl2pt* or *egl4e* fail to specify an oocyte (Mach et al., 1997; Navarro et al., 2004). Consequently, the effects of these mutations on mRNA localization could not be studied.

**Fig. 1:**
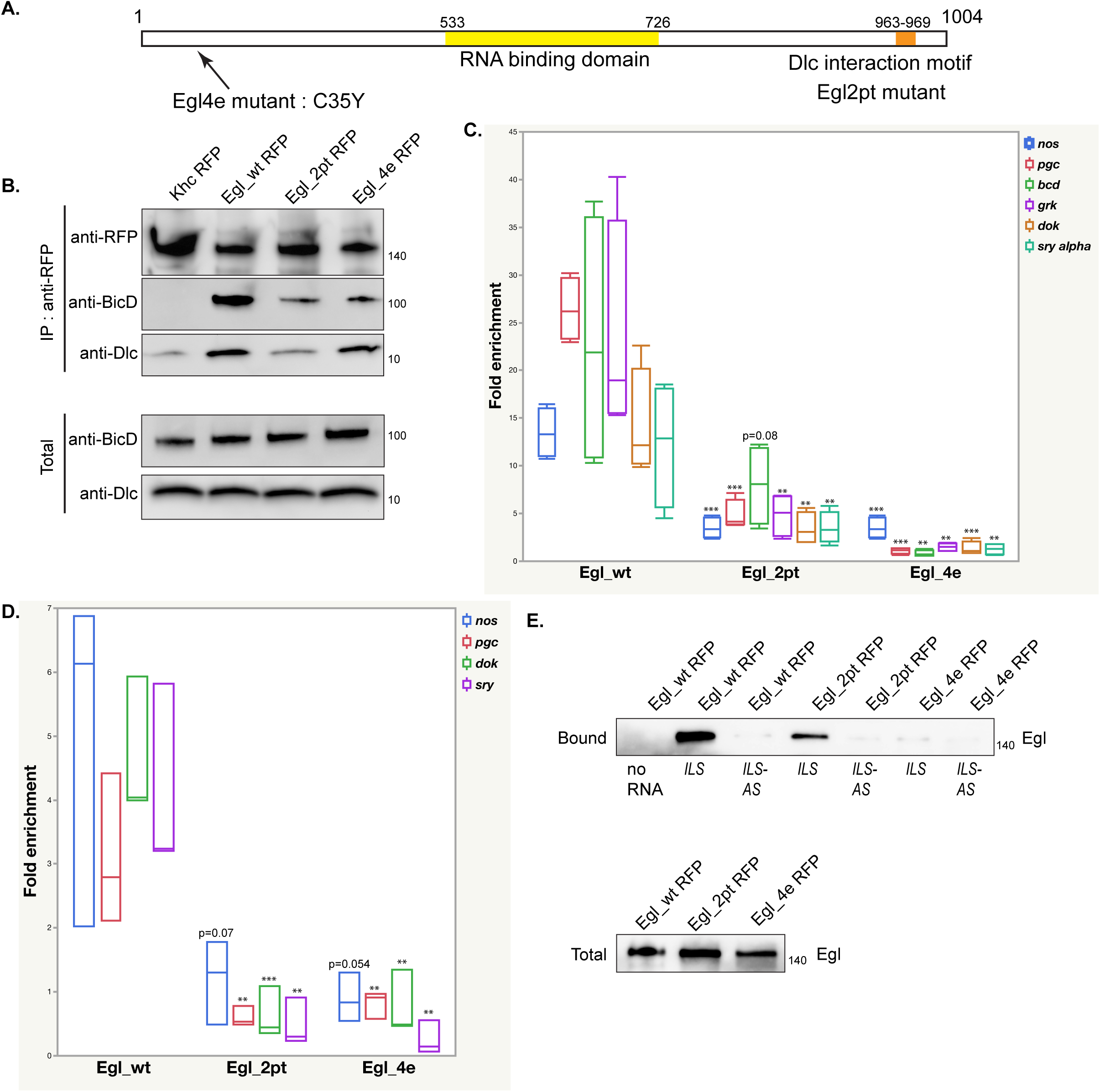
Egl_2pt and Egl_4e are deficient for binding RNA. **(A)** Schematic illustrating the domain structure of Egl. The RNA binding domain, the position of Egl_4e (C35Y) and Egl2pt (S965K, S969R) are indicated. **(B)** A co-immunoprecipitation was set up using ovarian lysates from strains expressing Khc-RFP, Egl_wt-RFP, Egl_2pt-RFP or Egl_4e-RFP. The strains expressing the Egl transgenes were also depleted of endogenous Egl. Lysates were incubated with RFP-Trap beads. After binding and wash steps, the bound proteins were eluted and analyzed by western blotting using the indicated antibodies. Wild-type Egl was able to co-precipitate BicD and Dlc. Egl_4e-RFP was able to co-precipitate Dlc but not BicD. Egl_2pt-RFP was deficient for binding both Dlc and BicD. **(C)** The same lysates were incubated with RFP-trap beads. After binding and wash steps, the bound RNA was extracted, reverse transcribed and analyzed using quantitative PCR. **(D)** A similar experiment to panel C was performed using lysates from 0 to 8 hour embryos. In comparison to wild-type, Egl_2pt and Egl_4e were defective for binding the indicated RNA cargoes in ovaries (C) and embryos (D). Panel C represents 4 biological replicates whereas Panel D represents 3 biological replicates. **(E)** *ILS* localization element or its anti-sense (*ILS-AS*) was bound to beads using a streptavidin aptamer. The beads were incubated with ovarian lysates from strains expressing Egl_wt-RFP, Egl_2pt-RFP or Egl_4e-RFP and depleted of endogenous Egl. After binding, RNA-associated proteins were eluted and analyzed by blotting using an RFP-antibody. A total fraction is also shown. Egl_2pt and Egl_4e are both defective for binding *ILS*. *** p<0.001, ** p <0.05 (Unpaired *t*-test).

In order to overcome this early requirement for Egl, we developed an shRNA depletion strategy (Sanghavi et al., 2016). By expressing an shRNA against *egl* using a stage-specific driver, we were able to generate egg chambers that were depleted of Egl within the germline. We reasoned that we could use a similar approach to address the functional role of the Egl/Dlc and Egl/BicD interactions. Transgenes were generated that encode either wild-type *egl*, *egl_2pt*, or *egl_4e*. The constructs were tagged on the C-terminus with RFP and contained silent mutations that made them refractory to the *egl* shRNA. All three transgenes were integrated at the same genomic locus and brought into the background of the *egl* shRNA strain. Thus, we were able to generate flies that were depleted of endogenous Egl, but expressed either wild-type or mutant versions of Egl in mid to late stage egg chambers and early embryos.

As expected, Egl_wt interacted efficiently with BicD and Dlc (Fig.1B). Also as expected, Egl_4e was able to bind Dlc but was defective for BicD binding (Fig.1B). Consistent with published results (Navarro et al., 2004), Egl_2pt was compromised for Dlc binding (Fig. 1B). Unexpectedly, this mutant was also defective for binding BicD (Fig.1B). A similar result was obtained by performing this experiment in *Drosophila* S2 cells (Supplemental Fig. 1A). Thus, loss of Dlc binding somehow also affects the Egl/BicD interaction.

We next examined the ability of these mutants to interact with native mRNAs. Ovarian or embryonic lysates were prepared from flies expressing wild-type or mutant RFP-tagged Egl. A strain expressing RFP tagged Kinesin heavy chain (Khc) was used as a negative control. Lysates were incubated with RFP-trap beads, the co-precipitating RNAs were extracted and analyzed by reverse transcription followed by quantitative PCR. *bicoid* (*bcd*) and *gurken* (*grk*) mRNAs are known targets of Egl (Bullock et al., 2001; Dienstbier et al., 2009), whereas *nanos* (*nos*), *polar granule component* (*pgc*), *dok* and *sry alpha* were recently identified as Egl cargoes using RIP-seq (Vazquez-Pianzola et al., 2017). In contrast to wild-type, Egl_2pt and Egl_4e were defective for binding mRNA cargo in ovaries and embryos (Fig.1C and D).

In order to validate this result, we examined the ability of wild-type or mutant Egl to bind to a localization element from the *I* factor retrotransposon (*ILS*). The anti-sense version of *ILS* (*ILS-AS*) was used as a control for specificity. *ILS* and *ILS-AS* containing a streptavidin binding apatamer were bound to beads as previously described (Dienstbier et al., 2009). Ovarian lysates were then incubated with the RNA-bound beads. Bound proteins were analyzed by western blotting. Consistent with results obtained using native mRNAs, wild-type Egl efficiently associated with *ILS*, but showed minimal binding to *ILS-AS* (Fig. 1E). By contrast, Egl_2pt displayed reduced binding to *ILS* yet was able to distinguish between *ILS* and *ILS-AS* (Fig. 1E). Egl_4e on the other hand, displayed greatly reduced binding to both *ILS* and *ILS-AS* (Fig. 1E).

Collectively, these results suggest that compromising the ability of Egl to interact with Dlc and BicD also affects its ability to bind RNA. Previous studies have shown that BicD is required for Egl to specifically bind RNA localization sequences (Dienstbier et al., 2009). Thus, the inability of Egl4e to bind RNA is not entirely surprising. However, a role for Dlc in mediating Egl mRNA binding has not been reported.

Although Dlc is a component of the Dynein motor, it also interacts with many proteins in a Dynein-independent manner (Rapali et al., 2011). The available evidence suggests that one of the main functions of Dlc is to facilitate dimerization or oligomerization of proteins (Barbar, 2008). In this regard, Egl has been shown to exist as a dimer in ovarian lysates (Mach et al., 1997). We therefore hypothesized that Dlc is required for Egl dimerization, and that dimerization is a pre-requisite for efficient RNA binding. In addition, although Egl and BicD can directly interact, the Bullock and Trybus labs recently demonstrated that this interaction is greatly enhanced by RNA (McClintock et al., 2018; Sladewski et al., 2018). Thus, we further posit that by affecting Egl’s ability to bind RNA, loss of dimerization also affects its ability to interact with BicD.

In order to test this hypothesis, we immunoprecipitated wild-type or mutant RFP-tagged Egl in strains that also expressed endogenous Egl. Consistent with previous results (Mach et al., 1997), Egl_wt-RFP was able to co-precipitate endogenous Egl (Fig. 2A). A similar result was obtained for Egl_4e-RFP. Consistent with our hypothesis, Egl_2pt-RFP was unable to co-precipitate endogenous Egl (Fig. 2A). In order to test dimerization more directly, we co-expressed GFP and FLAG versions of wild-type or mutant Egl in S2 cells. Whereas wild-type Egl-GFP was able to co-precipitate wild-type Egl-FLAG, Egl_2pt-GFP was unable to co-precipitate Egl_2pt-FLAG (Fig. 2B). We therefore conclude that Egl_2pt is dimerization defective. We further demonstrate that Egl dimerization is independent of its of RNA binding activity. An Egl construct lacking the RNA binding domain was able to dimerize and interact with Dlc (Supplemental Fig. 1B and Fig. 2C). As expected given recent findings (McClintock et al., 2018; Sladewski et al., 2018), loss of RNA binding significantly compromised the Egl-BicD interaction (Fig. 2C).

**Fig. 2:**
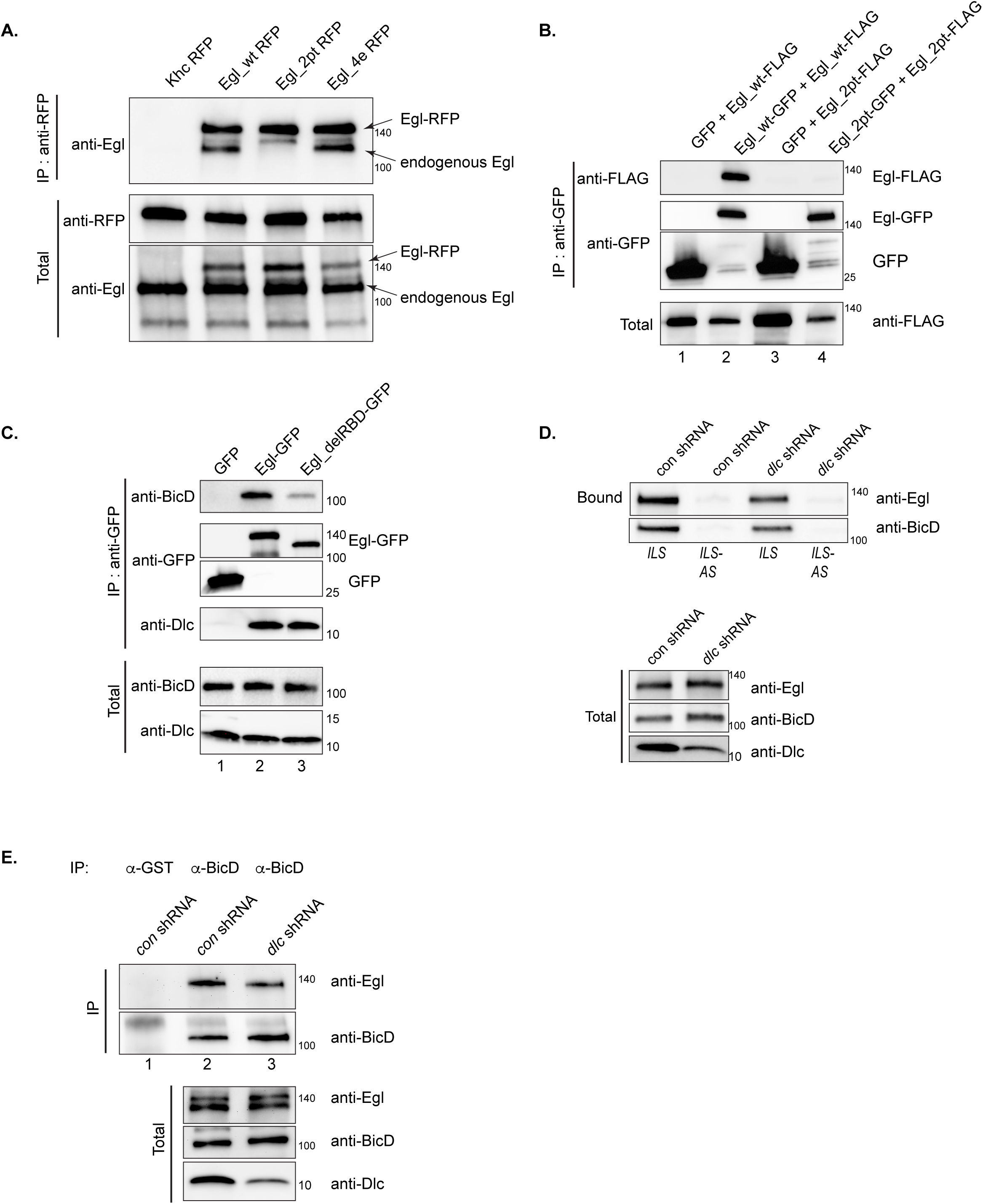
Dlc is required for Egl dimerization. **(A)** Ovarian lysates were prepared from strains expressing Khc-RFP, Egl_wt-RFP, Egl_2pt-RFP or Egl_4e-RFP. The strains also expressed endogenous Egl. The RFP tagged proteins were immunoprecipitated using RFP-trap beads and the co-precipitating proteins were analyzed by western blotting using the indicated antibodies. The total fraction is shown. Egl_wt-RFP and Egl_4e-RFP were able to co-precipitate endogenous Egl, whereas Khc-RFP and Egl_2pt-RFP were not. **(B)** A co-precipitation experiment was carried out using S2 cells transfected with the following constructs; lane 1 (GFP and Egl_wt-FLAG); lane 2 (Egl_wt-GFP and Egl_wt-FLAG), lane 3 (GFP and Egl_2pt-FLAG) and lane 4 (Egl_2pt-GFP and Egl_2pt-FLAG). The lysates were precipitated using GFP-Trap beads and the co-precipitated proteins were analyzed using the indicated antibodies. The total fraction is shown. Egl_2pt is dimerization defective. **(C)** A similar experiment was set up as in panel B using the following constructs; lane 1 (GFP), lane 2 (Egl_wt-GFP) and lane 3 (Egl_delRBD-GFP, an Egl construct lacking the RNA binding domain). Bound and total fractions are shown. Deletion of the RNA binding domain compromises the Egl-BicD interaction but not the Egl-Dlc interaction. **(D)** *ILS* or its anti-sense (*ILS-AS*) were bound to beads. Ovarian lysates were prepared from strains expressing a control shRNA (against the *white* gene) or an shRNA against *dlc*. These lysates were incubated with RNA-bound beads. After incubation, the bound proteins were eluted and analyzed by \ blotting using the indicated antibodies. The total fraction is also shown. Depletion of Dlc compromises the ability of Egl and BicD to associate with *ILS*. **(E)** The same lysates were used in a co-immunoprecipitation experiment using either an antibody against GST (lane 1) or BicD (lane 2 and 3). The bound proteins were analyzed using the indicated antibodies. Bound and total fractions are shown. Depletion of Dlc reduces the Egl-BicD interaction.

The above experiments involve a mutant of Egl that is defective for Dlc binding. If our hypothesis is correct, we should obtain similar results by depleting Dlc. Genetic loss of Dlc is lethal and germline clones do not specify an oocyte (Dick et al., 1996; Navarro et al., 2004). We therefore attempted to deplete Dlc using a specific shRNA and the same driver that was successful in depleting Egl (Sanghavi et al., 2016). Unfortunately this resulted in an oogenesis block (Supplemental Fig. 1C). We therefore used a different maternal driver that is expressed in slightly later stage egg chambers (Sanghavi et al., 2016). Although Dlc could be depleted using this driver, the knock-down was incomplete (Fig. 2D). Consistent with our hypothesis, depletion of Dlc partially compromised the ability of Egl to interact with RNA and BicD (Fig. 2D, E). The binding deficit observed with this strategy is not as strong as that observed in the Egl_2pt background. We attribute this difference to the incomplete depletion of Dlc using this driver and shRNA combination.

These findings raise an important question. Is the role of Dlc in this pathway simply to mediate Egl dimerization? If this is true, we might be able to bypass the function of Dlc by artificially dimerizing Egl. Sladewski and colleagues used a leucine zipper to artificially dimerize a truncated version of Egl (Sladewski et al., 2018). Using a similar approach, we observed that although Egl_2pt does not dimerize, Egl_2pt containing a C-terminal leucine zipper efficiently dimerizes (Fig. 3A). As expected, Egl_2pt-zipper was still unable to bind Dlc. However, despite this inability, Egl_2pt-zipper was able to interact with BicD (Fig. 3B). Importantly, in contrast to Egl_2pt, Egl_2pt-zipper was able to bind *ILS* and *TLS* (the localization element from *K10* mRNA), but displayed minimal binding to *ILS-AS* (Fig. 3C, D). Thus, RNA binding activity is also restored upon artificial dimerization of Egl. Based on these results we conclude that the primary function of Dlc in this pathway is to mediate Egl dimerization. Once dimerization is restored, RNA and BicD binding are also restored.

**Fig. 3:**
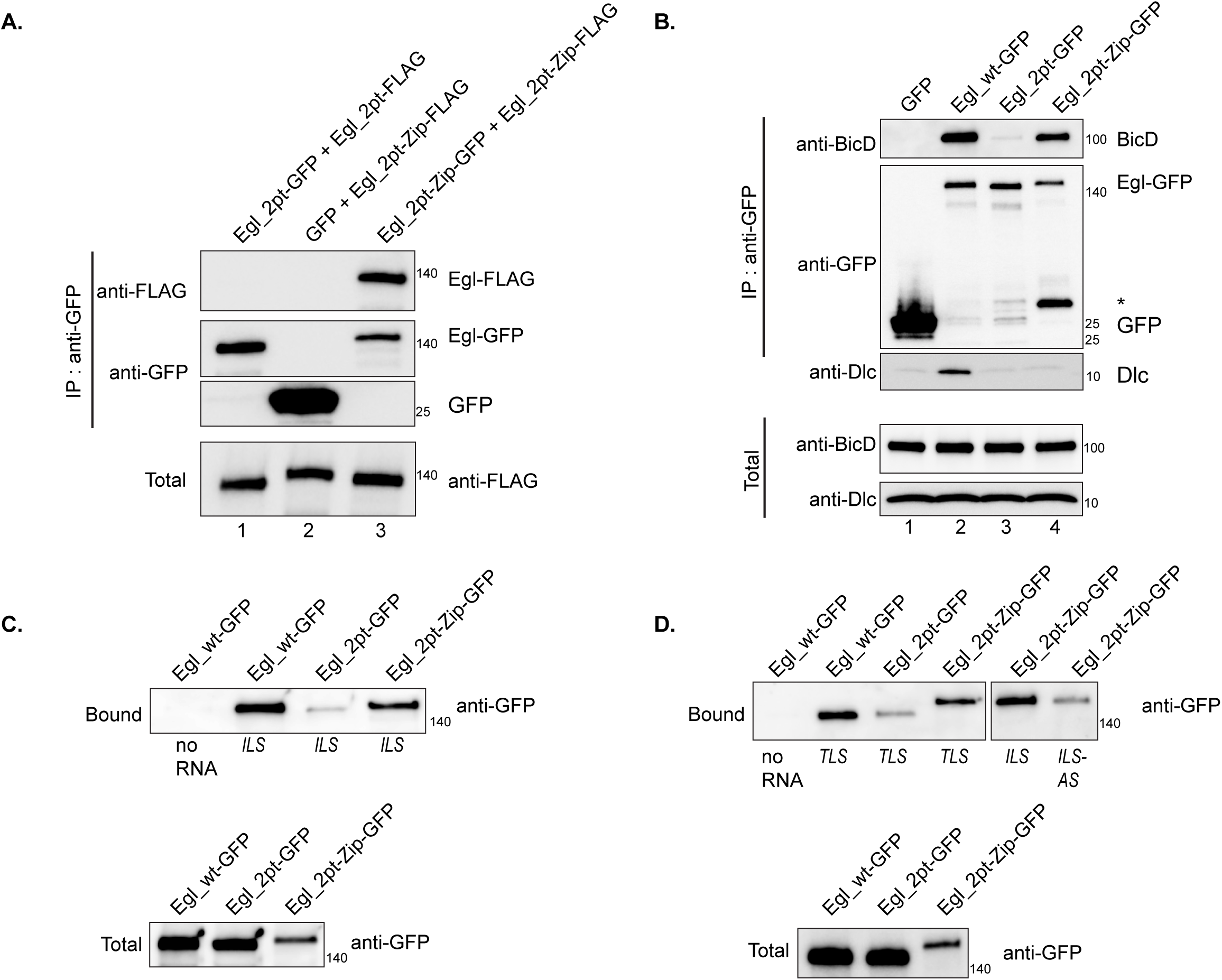
Artificial dimerization of Egl_2pt restores RNA and BicD binding. **(A)** A co-precipitation experiment was carried out using S2 cells transfected with the following; lane 1 (Egl_2pt-GFP and Egl_2pt-FLAG), lane 2 (GFP and Egl_2pt-Zip-FLAG), and lane 3 (Egl_2pt-Zip-GFP and Egl_2pt-Zip-FLAG). Zip refers to a leucine zipper motif. The lysates were incubated with GFP-Trap beads. The co-precipitating proteins were analyzed using the indicated antibodies. Insertion of the leucine zipper restores Egl_2pt dimerization. **(B)** A similar experiment was set up using the following constructs; lane 1 (GFP), lane 2 (Egl_wt-GFP), lane 3 (Egl_2pt-GFP) and lane 4 (Egl_2pt-Zip-GFP). After incubation with GFP-trap beads, the co-precipitating proteins were analyzed by western blotting. Egl_2pt-Zip-GFP is able to associate with BicD. *a degradation product is consistently seen in the Egl_2pt-Zip-GFP lane. **(C)** *ILS* was bound to beads and incubated with S2 cell lysates expressing the indicated constructs. **(D)** The *TLS* localization element, *ILS* or its anti-sense (*ILS-AS*) were bound to beads as indicated. The RNA-bound beads were incubated with S2 cells lysates expressing the indicated constructs. The bound proteins were analyzed by blotting. Although Egl_2pt is compromised for RNA binding, artificial dimerization (Egl_2pt-Zip) restores RNA binding.

We next examined the localization of Egl, Dynein, and BicD. Wild-type Egl, Dynein heavy chain (Dhc), the motor component of the Dynein motor, and BicD were all enriched in the oocyte of early stage egg chambers (Fig. 4A, G, Supplemental Fig. 2A) (Li et al., 1994; Mach et al., 1997). By contrast, Egl_2pt and Egl_4e were diffusely distributed with little oocyte enrichment (Fig. 4C, E, G). A similar, albeit weaker phenotype was observed for Dhc and BicD in these egg chambers (Fig. 4C’, E’, G, Supplemental Fig. 2B, C). The oocyte localization of BicD depends on Egl (Mach et al., 1997). Thus, one explanation for the weaker localization defect observed for Dhc and BicD in these mutant backgrounds is that at this stage of development endogenous Egl is not yet completely depleted (Sanghavi et al., 2016). As such, the partial oocyte enrichment of Dhc and BicD in these mutants is likely to be mediated by this residual endogenous Egl.

**Fig. 4:**
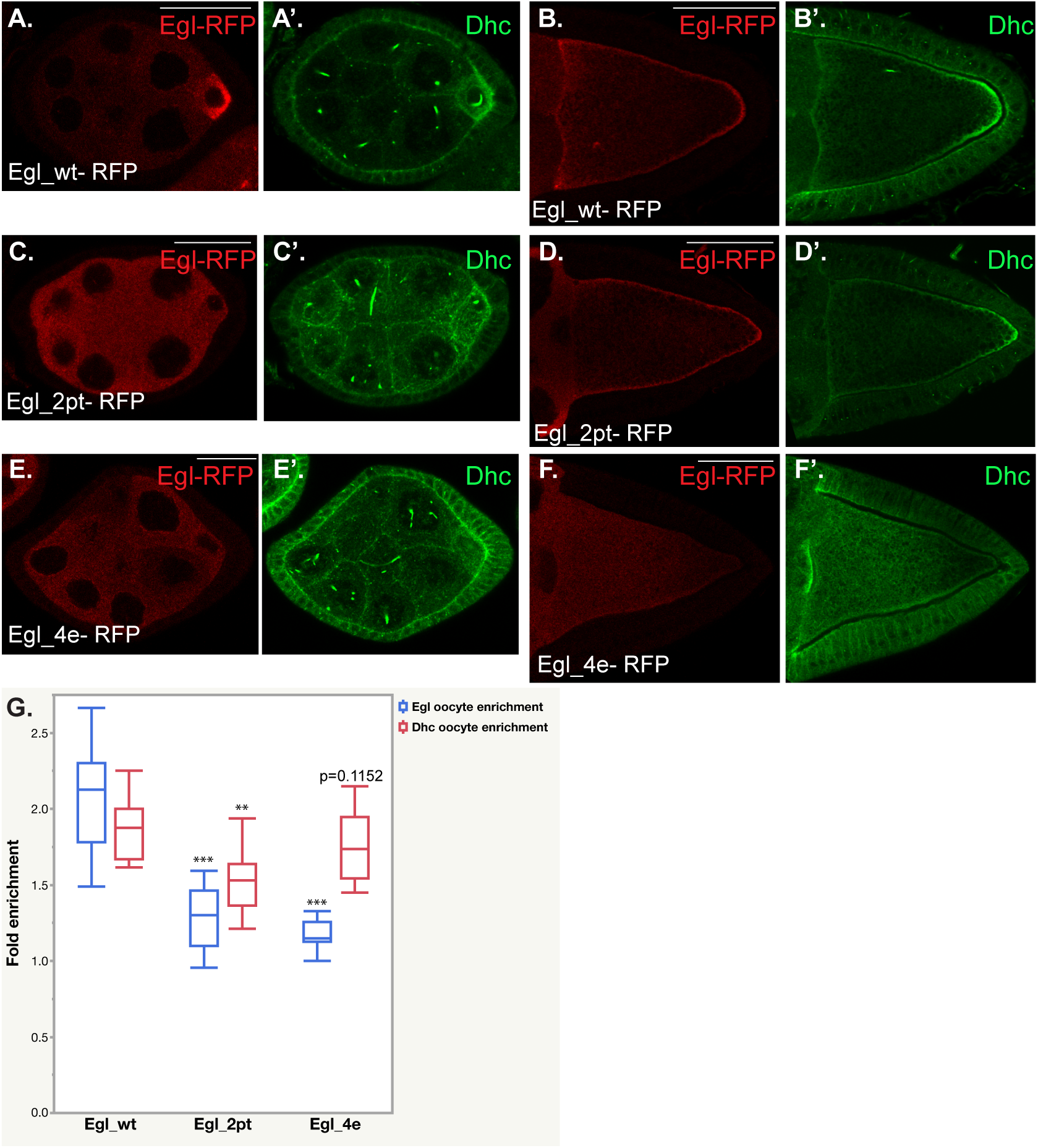
Localization of Egl and Dynein. **(A-F)** Ovaries from strains expressing Egl_wt-RFP (A, B), Egl_2pt-RFP (C, D) or Egl_4e-RFP (E, F) and depleted of endogenous Egl were fixed and processed using antibodies against RFP (A-F, red) and Dynein heavy chain (Dhc, A’-F’, green). Stage 5 (A, C, E) and stage 10 (B, D, F) egg chambers are depicted. **(G)** The oocyte enrichment of Egl and Dhc in stage 5 egg chambers was quantified. n=15, *** p<0.001, ** p <0.05 (Unpaired *t*-test). The scale bar in A, C and E is 25 microns; the scale bar in B, D, and F is 50 microns.

In stage 10 egg chambers expressing wild-type Egl, Dhc was partially enriched at the posterior pole, whereas Egl and BicD were localized to the oocyte cortex (Fig. 4B, Supplemental Fig. 2D). A similar pattern was observed in egg chambers expressing Egl_2pt (Fig. 4D, Supplemental Fig. 2E). Thus, although the transport of these factors is defective in early stage Egl_2pt egg chambers, they are restored to a normal pattern by stage 10. By contrast, all three proteins were significantly delocalized in stage 10 egg chambers expressing Egl_4e (Fig.4F, Supplemental Fig. 2F).

Knock down of Egl results in profound defects in the organization of oocyte microtubules (Sanghavi et al., 2016). We therefore examined microtubule organization in strains expressing Egl_2pt and Egl_4e. Kinesin β-gal localization was used to reveal the distribution of microtubule plus-ends, whereas gamma tubulin serves a marker for minus-ends (Clark et al., 1994; Cha et al., 2002). This analysis revealed that whereas plus and minus ends were correctly localized in egg chambers expressing wild-type Egl and Egl_2pt, both markers were delocalized in mutants expressing Egl_4e (Fig.5 A-F). The oocyte nucleus, which localizes to the dorsal-anterior region in a microtubule-dependent manner (Zhao et al., 2012), was correctly localized in strains expressing wild-type Egl and Egl_2pt but was sometimes delocalized in egg chambers expressing Egl_4e (Fig. 5G-I). Thus, microtubule organization is preserved in Egl_2pt mutants but is disrupted in Egl_4e mutants. Given that both mutants are deficient for binding BicD and RNA, we do not have an explanation for why microtubules are disorganized in the Egl_4e background.

**Fig. 5:**
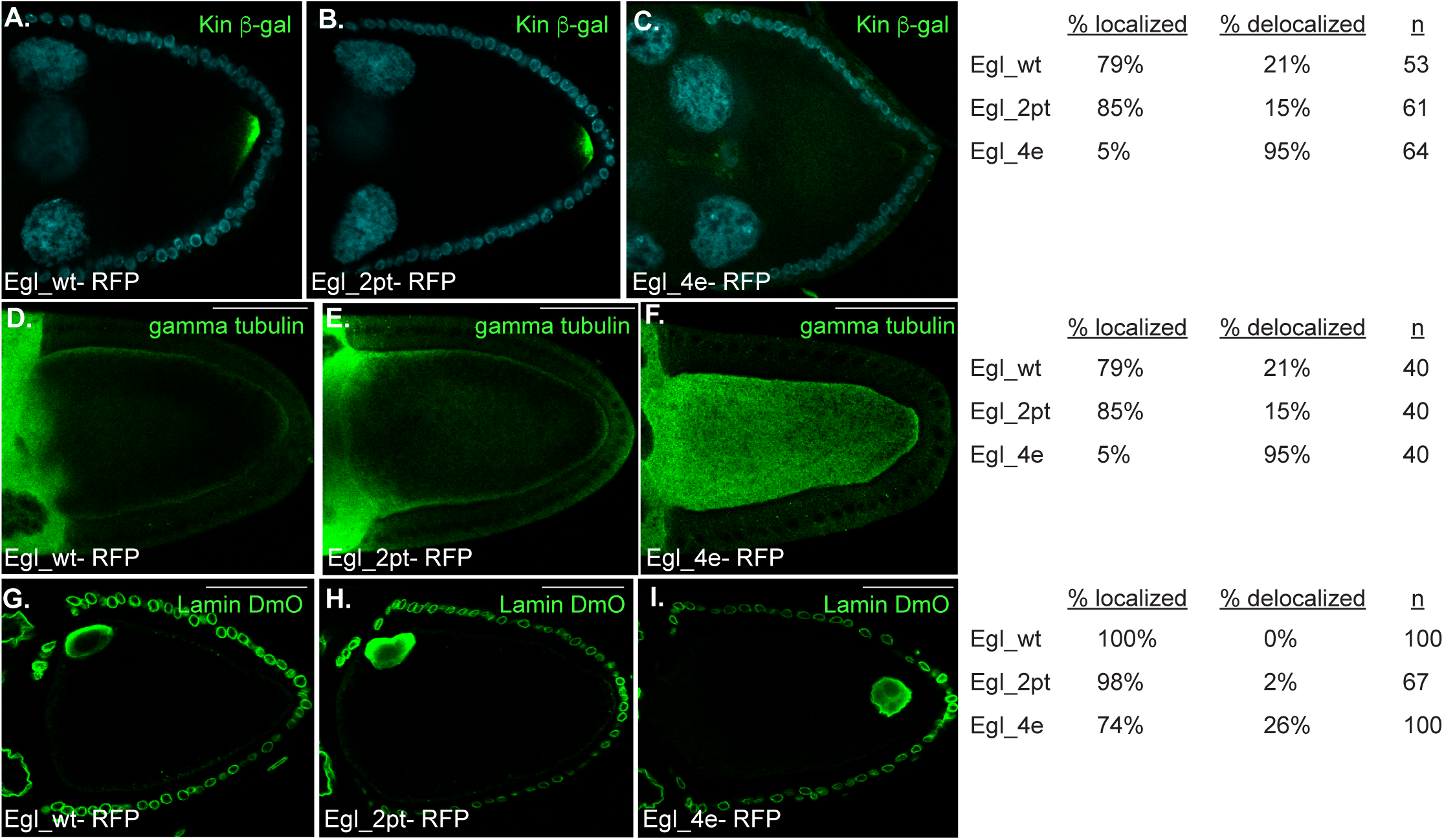
Microtubules are disorganized in Egl_4e mutants. **(A-C)** Strains co-expressing Egl_wt-RFP, Egl_2pt-RFP, or Egl_4e-RFP and Kinesin β-gal were fixed and processed for immunofluorescence using an antibody against beta-galactosidase (green). Phenotype quantifications are indicated. **(D-I)** Ovaries from strains expressing Egl_wt-RFP, Egl_2pt-RFP, or Egl_4e-RFP and depleted of endogenous Egl were fixed and processed for immunofluorescence using an antibody against gamma tubulin (D-LF) and Lamin DmO (G-I). Phenotype quantifications are indicated.

We next examined mRNA localization. The localization of *oskar* (*osk*) mRNA at the posterior of stage 10 egg chambers (Ephrussi et al., 1991; Kim-Ha et al., 1991) was maintained in wild-type Egl and Egl_2pt mutants. However, *osk* was significantly delocalized in mutants expressing Egl_4e (Fig. 6A-C). The delocalization of *osk* mRNA in Egl_4e mutants likely results from the disorganized microtubule cytoskeleton present in these egg chambers.

**Fig. 6:**
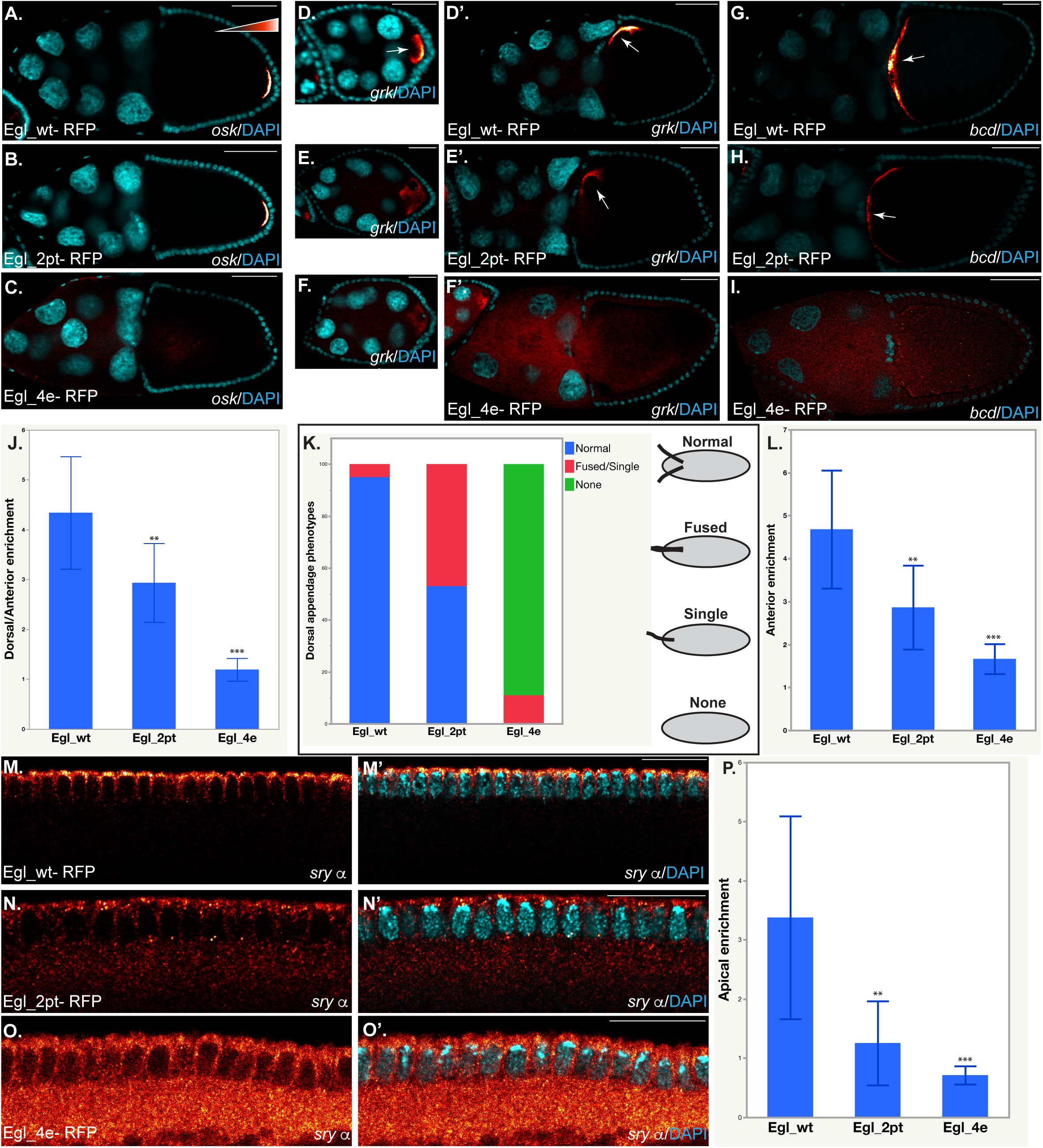
RNA localization in Egl wild-type and mutant backgrounds. **(A-I).** Ovaries from strains expressing Egl_wt-RFP, Egl_2pt-RFP, or Egl_4e-RFP and depleted of endogenous Egl were fixed and processed for single molecule FISH (smFISH) using probes against *osk* (A-C), *grk* (D-F) or *bcd* mRNA (G-I). **(J)** The dorsal-anterior enrichment of *grk* mRNA was quantified in strains expressing Egl_wt-RFP, Egl_2pt-RFP and Egl_4e-RFP; n=10. **(K)** The dorsal appendage phenotype in mature eggs from these strains was scored; n=100. **(L)** The anterior enrichment of *bcd* in these strains was quantified; n=10. **(M-O)** Embryos from strains expressing Egl_wt-RFP, Egl_2pt-RFP and Egl_4e-RFP were fixed and processed for smFISH using probes against *sry alpha*. **(P)** The apical enrichment of sry alpha in these embryos was quantified, n=10 for Egl_wt and Egl_2pt and n=7 for Egl_4e. *In situ* signal is depicted using a red to white LUT; red represent low-intensity signal and white represent high-intensity signal. *** p<0.001, ** p < 0.05 (Unpaired *t*-test). The scale bar for D, E and F is 25 micros. For the rest of the images, the scale bar is 50 microns.

*grk* mRNA is enriched in the oocyte of early stage egg chambers and is localized at the dorsal-anterior cortex by stage 10 (Neuman-Silberberg et al., 1993). As expected, given the compromised localization of Egl and Dhc in early stage egg chambers (Fig. 4C, E), the oocyte enrichment of *grk* was reduced in egg chambers expressing Egl_2pt and Egl_4e (Fig. 6D-F). Furthermore, in comparison to wild-type, the dorsal-anterior enrichment of *grk* mRNA was reduced in Egl_2pt mutants and was completely abrogated in Egl_4e egg chambers (Fig. 6D-F, J). Grk protein, produced at the dorsal corner of the oocyte, is secreted and signals the overlying follicle cells to adopt dorsal fate (Neuman-Silberberg et al., 1993). The end result is formation of dorsal appendages in mature eggs. Consistent with the mRNA localization result, normal dorsal appendage formation was reduced in Egl_2pt mutants and was severely compromised in Egl_4e flies (Fig. 6K).

*bcd* mRNA localizes at the anterior margin of stage 10 egg chambers where it persists until early stages of embryogenesis (Berleth et al., 1988). As observed for *grk*, the anterior enrichment of *bcd* mRNA was reduced in Egl_2pt mutants and virtually eliminated in Egl_4e mutants (Fig. 6G-I, L).

Finally, we examined the localization of *dok* and *sry alpha* in oocytes and embryos. These mRNAs were identified as Egl cargoes by the Suter lab and our results are consistent with their findings (Vazquez-Pianzola et al., 2017) (Fig. 1C, D). Both *dok* and *sry alpha* are enriched within the oocyte of early stage wild-type egg chambers (Vazquez-Pianzola et al., 2017) (Supplemental Fig. 2G, J). This localization pattern was compromised in egg chambers expressing Egl_2pt or Egl_4e (Supplemental Fig. 2H-L). In addition to the oocyte, both mRNAs are also localized to the apical surface of blastoderm embryos (Vazquez-Pianzola et al., 2017). Our probe set against *dok* revealed an apical enrichment pattern in only a subset of blastoderm embryos (data not shown). Therefore, we focused our analysis on the localization of *sry alpha*. Consistent with published results, *sry alpha* was strongly localized at the apical surface of embryos expressing wild-type Egl (Vazquez-Pianzola et al., 2017) (Fig. 6M, P). By contrast, this enrichment was greatly reduced in Egl_2pt mutants and was completely disrupted in embryos expressing Egl_4e (Fig. 6N-P).

The severe localization defects observed for *grk*, *bcd* and *sry alpha* in Egl_4e mutants are likely to be caused by the disorganized microtubule architecture present in these egg chambers.

Based on these results, we propose the following model (Fig. 7). Dlc is required for dimerization of Egl. Dimeric Egl is able to more efficiently associate with RNA cargo that is destined for localization. Cargo association in turn promotes the interaction of Egl with BicD, an adaptor for the Dynein motor. As such, disrupting the Egl/Dlc interaction results in mRNA being inefficiently associated with Dynein, and therefore incompletely localized within the cell.

**Fig. 7:**
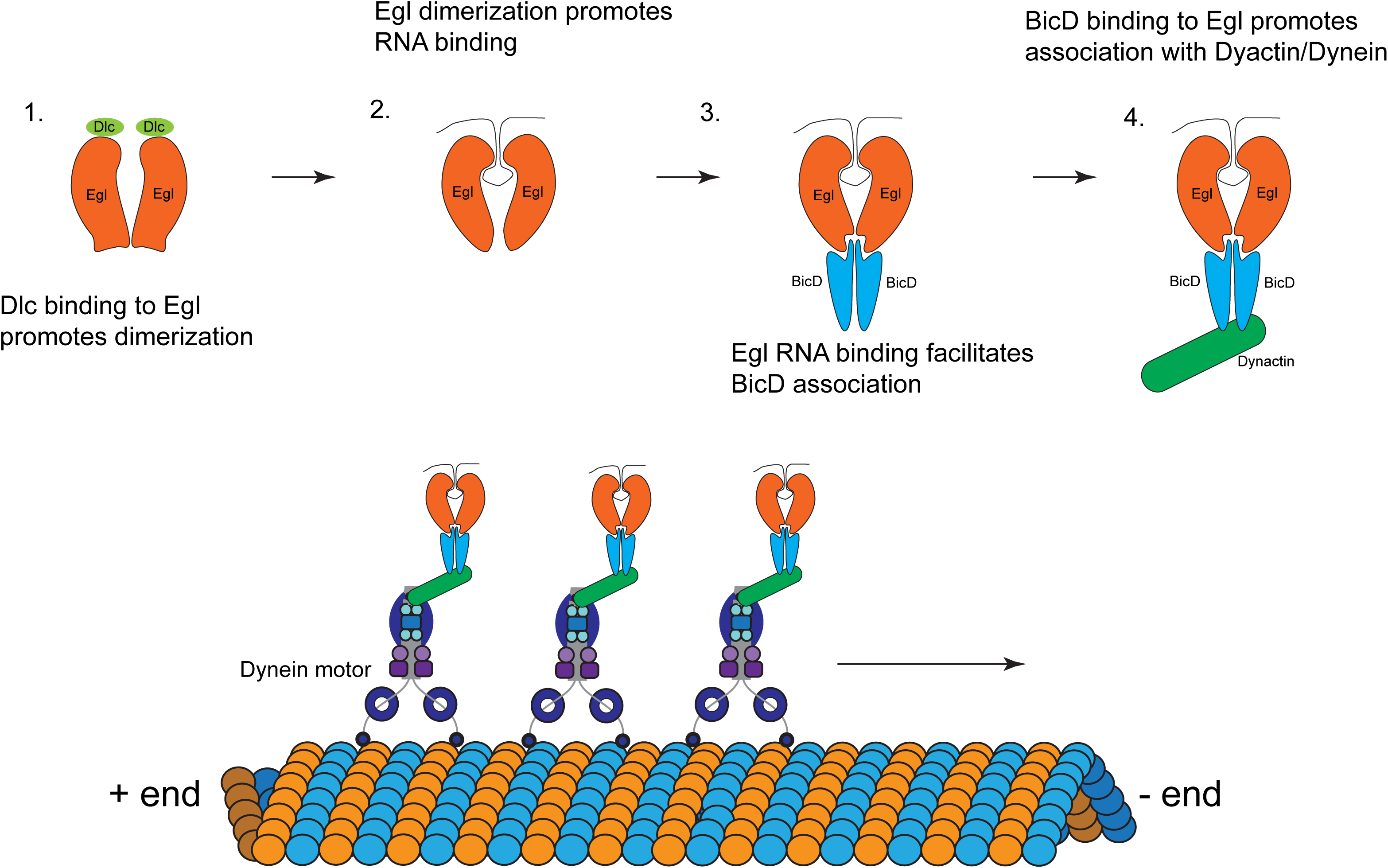
Model. A model to illustrate the mechanism by which Dlc and BicD function to link the Egl/mRNA complex to the Dynein motor.

## Discussion

Egl has been known to physically interact with Dlc for many years (Navarro et al., 2004). However, the functional significance of this interaction has remained unknown. The reason for this has to do with the fact that Egl and Dlc are both required for specification of the oocyte. Genetic loss of either of these gene products arrests oogenesis at very early stages (Theurkauf et al., 1993; Carpenter, 1994; Navarro et al., 2004). As such, classical genetic approaches could not be used to study mRNA localization in these mutant backgrounds.

In a recent study, we described a strategy to overcome this limitation with respect to Egl (Sanghavi et al., 2016). By using shRNA mediated depletion and a germline specific driver that is only expressed after oocyte specification, we were able to generate mid and late stage egg chambers that were depleted of endogenous Egl. This analysis revealed that Egl was required for the localization of *osk*, *bcd* and *grk* mRNAs in the *Drosophila* oocyte (Sanghavi et al., 2016). In this report, we used a similar strategy to examine the functional role of the Egl/Dlc interaction in oocytes and embryos.

Disrupting the Egl/Dlc interaction greatly diminishes the ability of Egl to associate with mRNA localization sequences (Fig. 1). On a mechanistic level, our findings indicate that Dlc is primarily required for Egl dimerization and that Egl dimerization is linked to its RNA binding activity (Fig. 2). If Egl is unable to dimerize, it binds RNA localization sequences much less efficiently; its specificity for localization sequences, however, is unaffected. Consistent with our hypothesis, artificial dimerization of the Egl_2pt mutant restored RNA binding, and consequently, the ability of this mutant to interact with BicD (Fig. 3).

The net result of disrupting the Egl/Dlc interaction is defective mRNA localization (Fig. 6). *grk*, *bcd*, and *sry alpha* were inefficiently localized in Egl_2pt mutants. It is likely that the same phenotype would also be observed for other Dhc-localized mRNA cargo. Although *grk*, *bcd* and *sry alpha* were not efficiently localized in *egl_2pt* mutants, they were not completely delocalized either. This suggests that the residual RNA binding activity of Egl_2pt might be sufficient to couple a certain fraction of these cargo molecules to Dynein. However, this might not be true for all cargoes. For instance, cargoes localized during early stages of egg chamber development might be particular sensitive to disruption of the Egl/Dlc interaction. In early-stage egg chambers, the oocyte enrichment of Egl_2pt is significantly compromised (Fig. 4C). Thus, cargoes localized during this time frame will likely not be correctly sorted. In this regard, it is interesting to note that flies homozygous for *egl_2pt* do not form mature eggs (Navarro et al., 2004). Rather, maturation is halted and oocyte fate is not maintained. It is possible that certain essential cargoes that are transported by Egl and Dynein are not correctly localized in this background, resulting in an oogenesis block. Additional studies might reveal the identity of these cargoes.

Another class of cargo that might be highly sensitive to disruption of the Egl/Dlc interaction are cargoes that not actively anchored at their site of localization. Both *grk* and *bcd* mRNAs have well-defined anchoring mechanisms (Delanoue et al., 2007; Weil et al., 2008; Trovisco et al., 2016). Thus, in the case of these mRNAs, even if transport is not that efficient, the presence of an anchoring mechanism can lessen the severity of the localization defect. Cargoes without such and anchoring mechanism would be significantly more delocalized.

Using purified proteins, the Bullock and Trybus labs recently demonstrated that a minimal complex consisting of an RNA-localization element, Egl, BicD, Dynactin and Dynein was sufficient for processive minus-end movement (McClintock et al., 2018; Sladewski et al., 2018). These studies also indicated that the presence of RNA stimulated the Egl-BicD interaction and that the majority of fast-moving complexes contained two copies of Egl (McClintock et al., 2018; Sladewski et al., 2018). Our results are consistent with these findings and indicate that prior to binding RNA and BicD, Egl is first dimerized with the help of Dlc. Our results also indicate that disrupting the Egl/Dlc interaction affects formation of this minimal transport complex.

The main difference between our findings and the conclusions drawn by the aforementioned in vitro studies has to do with the role of Dlc in RNA transport. Using purified complexes containing either wild-type or Egl_2pt (McClintock et al., 2018) or using Dynein complexes lacking Dlc (Sladewski et al., 2018), the authors conclude that Dlc is neither required for linking the Egl/RNA complex with Dynein, nor is it required for transport of said cargo. Although true under in vitro conditions using purified proteins, the situation in vivo, as reflected by our findings, indicates an important role for Dlc in this process. In the complex environment of the cell, Dlc is required for Egl dimerization and consequently for RNA binding. Thus, although Dlc is not required for linking the Egl/ RNA complex with Dynein, it is required at an earlier step; formation of a localization-competent transport particle.

## Acknowledgements

We thank Ruth Lehmann for providing the Egl antibody and Simon Bullock for providing the *ILS*, *ILS-AS* and *TLS* aptamer constructs and for helpful advise in performing the RNA binding experiments. We thank the Bloomington Stock Center, Developmental Studies Hybridoma Bank, and the *Drosophila* Genomics Resource Center for providing fly strains, antibodies, cell lines, and DNA constructs. This work was supported by a grant from the National Institutes of Health to G.B.G (R01GM100088).

## Materials and Methods

### Fly stocks

The following stocks were used: Oregon-R-C and w[1118] were used as wild-type (Bloomington stock center; #5 and #5905 respectively). The shRNA strains were: *eb1* shRNA (Bloomington stock center; #36680, donor TRiP); *white* shRNA (Bloomington stock center; #35573, donor TRiP); *egl* shRNA-1 (Bloomington stock center; #43550, donor TRiP); *ctp/lc8* shRNA (Bloomington stock center; #42862, donor TRiP). shRNA expression was driven using either P{w[+mC]=matalpha4-GAL-VP16}V37 (Bloomington Stock Center, #7063; donor Andrea Brand) for early-stage expression or w[*]; P{w[+mC]=matalpha4-GAL-VP16}V2H (Bloomington Stock Center, #7062; donor Andrea Brand) for mid-stage expression. Egl_wt-RFP, Egl_2pt-RFP, Egl_4e-RFP and Khc-RFP transgenes were constructed by cloning the respective coding regions into the pattB vector (Bischof et al., 2007). All four alleles were inserted at the attP1 site (Bloomington Stock Center, #34760) (Ni et al., 2009). The transgenic strains were injected by BestGene Inc. Microtubule plus-ends were marked using the Kin:βgal strain (Clark et al., 1994). Fly stocks and crosses used for these experiments were maintained at 25^0^C.

### Antibodies

The following antibodies were used for immunofluorescence: rat anti-RFP (Chromotek 1:1000); mouse anti-Dhc (Developmental Studies Hybridoma Bank, 1:250; donor J. Scholey); mouse anti-BicD (Clones 1B11 and 4C2, Developmental Studies Hybridoma Bank, 1:300; donor R. Steward); mouse anti-β-galactosidase (Promega, 1:2000); mouse anti-γ-tubulin (Sigma, 1:100); mouse anti-LaminDmO (Developmental Studies Hybridoma Bank, clone ADL84.12; 1:200; donor P. A. Fisher); goat anti-rat Alexa 594 (Life Technologies 1:400) goat anti-rabbit Alexa 594 and 488 (Life Technologies, 1:400 and 1:200 respectively); goat anti-mouse Alexa 594 and 488 (Life Technologies, 1:400 and 1: 200 respectively). The following antibodies were used for western blotting: rabbit anti-Egl (1:5000, from R. Lehmann); rabbit anti-Ctp (Abcam, 1:5000); mouse anti-GFP (Clontech, 1:5000) rat anti-RFP (Chromotek, 1:4000); mouse anti-BicD (Clones 1B11 and 4C2, Developmental Studies Hybridoma Bank, 1:300 each); anti-FLAG (Sigma Aldrich, 1:5000); goat anti-mouse HRP (Pierce, 1:5000); goat anti-rabbit HRP (Pierce, 1:5000); goat anti-rat HRP (Pierce, 1:5000). The following antibodies were used for immunoprecipitation: GST (Santa Cruz Biotechnology, 1:15) and mouse anti-BicD (Developmental Studies Hybridoma Bank; 1:30).

### DNA constructs

The transgenes for expressing Egl_wt, Egl_2pt, Egl_4e and Khc in the female germline were constructed by cloning the respective cDNAs into the pAttB vector (Bischof et al., 2007). The mutations were made using the Q5 site directed mutagenesis kit (New England Biolabs). The constructs contained the promoter from the maternal alpha tubulin gene, a C-terminal RFP tag, and the 3’UTR from the vasa gene. Two silent mutations were introduced into the constructs in order to make them refractory to the *egl* shRNA. In addition, a flexible linker was present in between the gene of interest and the RFP tag. The constructs were engineered using Gibson assembly (New England Biolabs). For expression in S2 cells, the same constructs were cloned into the pUASp-attB-K10 vector (Bischof et al., 2007) upstream of either a GFP, RFP, or 3xFLAG tag. The constructs were expressed in S2 cells by co-transfecting along with an Act5c-Gal4 plasmid. The leucine zipper from GCN4 (AAL09032.1) along with a preceding flexible linker sequence was synthesized with *Drosophila* codon optimization by Genewiz.

### Cell culture

The S2 cells used in this study were obtained from the *Drosophila* Genomics Resource Center. The cells correspond to the S2-DRSC line (stock number 181). The cells were grown in Schneider’s medium containing 10% heat inactivated fetal calf serum. S2 cells were transfected using Effectene (Qiagen) accord to the instructions provided by the manufacturer.

### Analyzing protein-protein interaction

Ovaries from the indicated genotypes were dissected and flash frozen in liquid nitrogen until ready to use. The ovaries were homogenized into lysis buffer (50mM Tris pH 7.5, 50mM NaCl, 0.2mM EDTA, 0.05% NP40 and Halt protease inhibitor cocktail, Pierce) and cleared by centrifugation at 10,000g at 4^0^C for 5min. Between 800ug and 1000ug total protein was used per immunoprecipitation. For ovarian lysates, immunoprecipitation was performed by incubating the lysates at 4^0^C for 1hour with RFP-trap beads (Chromotek). Subsequently, the beads were washed four times with wash buffer (50mM Tris pH 7.5, 200mM NaCl, 0.2mM EDTA, 0.05% NP40). The co-precipitating proteins were eluted in Laemmli buffer, run on a gel, and analyzed by western blotting. For examining protein-protein interaction using S2 cells, lysates were prepared using the indicated transfected cells. Lysis was performed using RIPA buffer (50mM Tris-Cl pH 7.5, 150nM NaCl, 1% NP40, 1mM EDTA). The lysates were cleared by centrifugation as described above and added to GFP-trap beads

### Analyzing in vivo protein-mRNA association

In order to examine binding to native mRNAs, ovaries were dissected and stored as described above. 0-8hr embryos were also collected, dechorinated, and flash frozen in liquid nitrogen. Between 800ug and 1000ug of ovarian or embryonic lysate was used per immunoprecipitation. The tagged proteins were immunoprecipitated using RFP-Trap beads using a previously described protocol (Sanghavi et al., 2016). The co-precipitating RNAs were reverse transcribed using random hexamers and Superscript III (Life Technologies) according to directions provided by the manufacturer. 167ng of cDNA was used in each quantitative PCR (qPCR) reaction (except for reactions with *grk* mRNA: 267ng) using the SsoAdvanced Universal SYBR Green Supermix (Bio-Rad). A Bio-Rad CFX96 Real-Time PCR System was used for this experiment. Fold enrichment was calculated by comparing ct values for each RNA to that obtained for γ-tubulin.

The following primers were used: *bcd* mRNA (5′-AATCGGATCAGCACAAGGAC-3′ and 5′-GCGTTGAATGACTCGCTGTA-3′), *grk* mRNA (5′-ATCCGATGGTGAACAACACA-3′ a n d 5 ′ - C G A C G A C A G C A T G A G G A G T A - 3 ′), *n o s* m R N A (5’ - TGTGGCGAAACATGTCGTAT-3’ and 5’-TCTCACAAAGACGCAGTGG-3’), *pgc* mRNA (5’-ACCCGAAAATGTGCGACTAC-3’ and 5’-ATCTCCATCTATCCGCGATG-3’), *sry-α* mRNA (5’GCAGTGAGCTGATTGCAGAG-3’ and 5’-AGGTGGGTGATGCAGGTTAC-3’), *d o k* m R N A (5’ - A T G T C C G C C A A G A T G A A G T C - 3’ a n d 5’ - GA T A TGCAGCTTTGCGTTGA-3’), and *γ-tubulin* mRNA (5 ′ - CCACCATCATGAGTCTGAGC-3′ and 5′-ACCGATGAGGTTGTTGTTCA-3′).

### In vitro protein/RNA interaction

*ILS*, *ILS-AS*, and *TLS* constructs containing a T7 transcription site and an aptamer with high affinity for streptavidin was obtained from Simon Bullock (Dix et al., 2013). The constructs were transcribed and purified as previously described (Dix et al., 2013). The RNA affinity purification was also performed as previously described with a few modifications (Dix et al., 2013). 5µg of RNA was first refolded in 10µL of *Drosophila* extraction buffer (DXB: 25mM HEPES pH 6.5, 50mM KCl, 1mM MgCl2, 250mM sucrose, 0.1% NP40, supplemented with 10µM MgATP and 1mM DTT at the time of use). Refolding was accomplished using a Bio-Rad C1000 Thermo Cycler with the following protocol: 56°C/5min, 37°C/10min, 25°C/10min. The refolded RNA was then incubated with High Capacity Streptavidin Agarose Beads (Pierce) in 90µL DXB for 1.5 hours at 4°C while nutating. Extracts were prepared from either frozen *Drosophila* ovaries or dechorionated embryos in lysis buffer (50mM Tris pH 7.5, 50mM NaCl, 0.2mM EDTA, 0.05% NP40 and Halt protease inhibitor cocktail, Pierce) using a sanitized pestle or in RIPA buffer (50mM Tris pH 7.5, 200mM NaCl, 0.2mM EDTA, 0.05% NP40) for S2 cells. Halt Protease Inhibitor mix (Thermo Scientific) and Rnase Out (Life Technologies) were added to the extracts. The beads were rinsed twice with the above lysis buffer prior to the addition of extracts. Typically between 800 to 1000ug of total protein was used in each binding experiment. The extracts were incubated with the RNA-bound streptavidin beads for 15 min at room temperature and a further 30 min at 4°C with nutation. The beads were then rinsed 4 times in lysis buffer, the bound proteins were eluted by boiling in Laemmli buffer and examined by western blotting.

### Immunofluorescence

Prior to dissection, flies were fattened on yeast pellets for 3 days. Ovaries were dissected as previously described (Liu et al., 2015) and fixed in 4% formaldehyde (Pierce) for 20 min at room temperature. The fixative was diluted in PBS. The immunofluorescence staining was performed as previously described (Liu et al., 2015).

The primary antibody was incubated in 1× PBST (PBS + 0.1% Triton X-100) + 0.2% BSA (Promega) overnight at 4°C. The next day, the samples were washed three times in PBST. The secondary antibody was diluted in 1× PBST + 0.2% BSA and incubated overnight at 4°C. The following day, the ovaries were washed three times in PBST, mounted on slides with Aqua-Poly/Mount (APM, Polysciences Inc.), and imaged.

### *In situ* hybridization

Prior to dissection, flies were fattened on yeast pellets for 3 days. Dissected ovaries and dechorionated embryos were fixed with 4% formaldehyde for 20 mins. Next, the fixative was removed and the ovaries were washed with PBST. Ovaries were further teased apart using a pipette. The ovaries or embryos were washed with 100% methanol for 5 min, then stored for at least 1 hour in 100% methanol at −20°C. The samples were then re-hydrated with three 10 min washes with a solution of PBST and 100% methanol (3:7, 1:1, 7:3) and rinsed four times with PBST. The samples were then washed for 10 minutes in Wash Buffer (4xSSC, 35% deionized formamide, 0.1% Tween-20). Fluorescent probes were diluted in Hybridization Buffer (10% dextran sulfate, 0.1mg/ml salmon sperm ssDNA, 100 µl vanadyl ribonucleoside (NEB biolabs), 20ug/ml RNAse-free BSA, 4x SSC, 0.1% Tween-20, 35% deionized formamide) and were applied overnight at 37°C. The following day, the samples were washed twice with pre-warmed Wash Buffer for 30 min. After two rinses with PBST and two rinses with PBS, the ovaries or embryos were mounted on slides using APM and imaged.

### Microscopy

Images were captured on a Zeiss LSM 780 upright microscope and processed for presentation using Fiji, Adobe Photoshop, and Adobe Illustrator. All imaging experiments were performed at the Augusta University Cell Imaging Core.

### Quantification

Quantifications of localized RNAs and proteins were carried out by measuring the average pixel intensity of the localized signal and dividing by the delocalized signal. For *grk* and *bcd*, the localized signal at the dorsal-anterior (*grk*) of the oocyte or at the anterior margin (*bcd*) of the oocyte was compared to the delocalized signal in the rest of the oocyte. For *sry alpha*, the localized signal above the blastoderm nuclei was compared to the delocalized signal below the nuclei. For quantifying Egl and Dhc oocyte enrichment, the signal in the oocyte was compared to the delocalized signal in the rest of the egg chamber. Quantifications were performed using Zeiss Zen Black software. An unpaired *t*-test was performed using standard deviation, mean, and the n value.

---

**Fig. S1:**
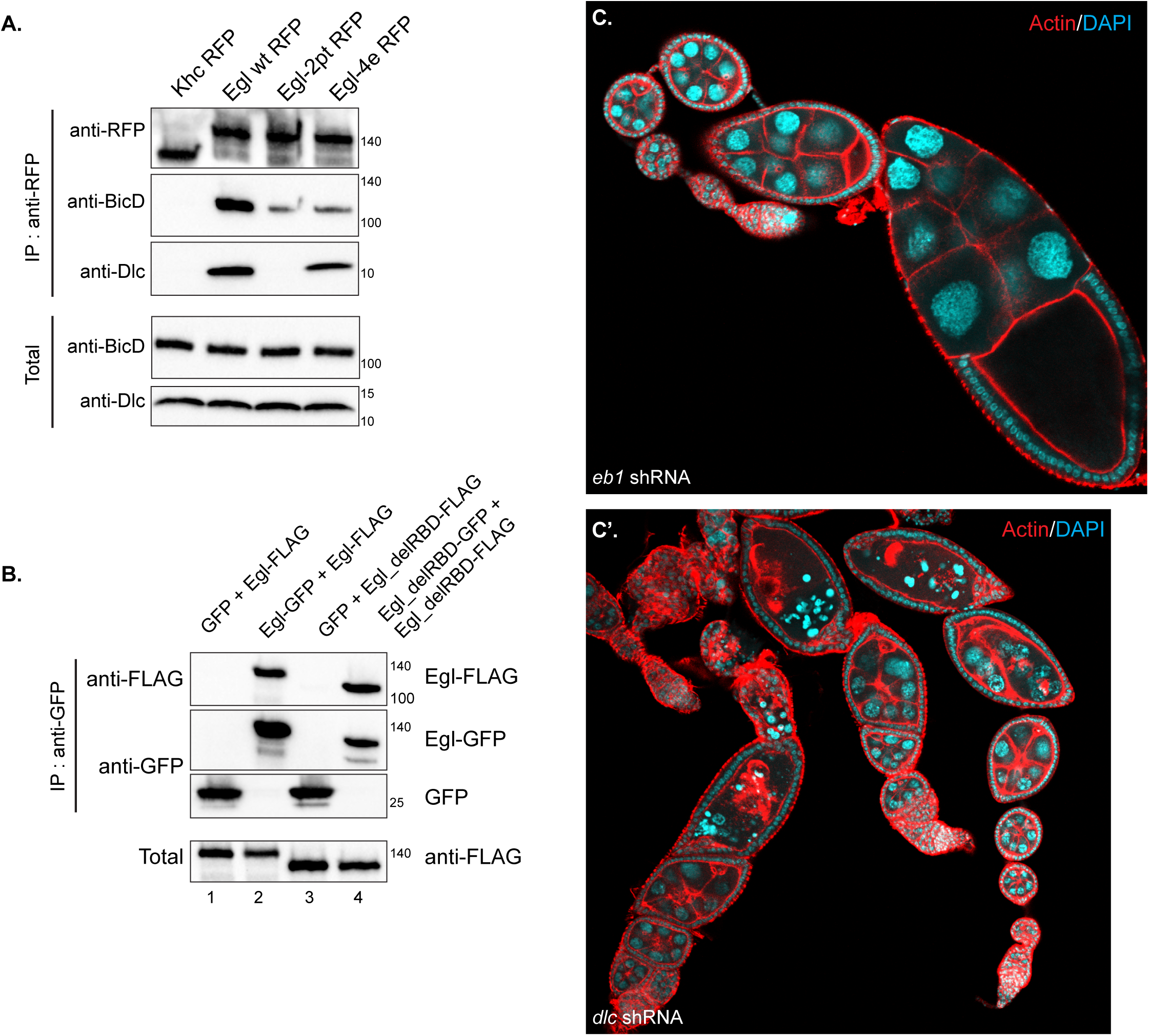
(A) A co-immunoprecipitation was set up using S2 cell lysates from strains expressing the indicated constructs. The lysates were incubated with RFP-Trap beads. After binding and wash steps, the bound proteins were eluted and analyzed by western blotting using the indicated antibodies. Wild-type Egl was able to co-precipitate BicD and Dlc. Egl_4e-RFP was able to co-precipitate Dlc but not BicD. Egl_2pt-RFP was deficient for binding both Dlc and BicD. (**B)** A co-immunoprecipitation experiment was set up using S2 cells expressing the indicated constructs. The lysates were incubated with GFP-trap beads. The bound proteins were analyzed using the indicated antibodies. **(C)** Ovaries from flies expressing either a control shRNA (*eb1* shRNA) or shRNA against *dlc* were fixed and processed using TRITC-Phalloidin and DAPI. The driver used for this experiment is restricted to the germline and is turned on early-stage egg chambers (Sanghavi et al., 2016).

**Fig. S2:**
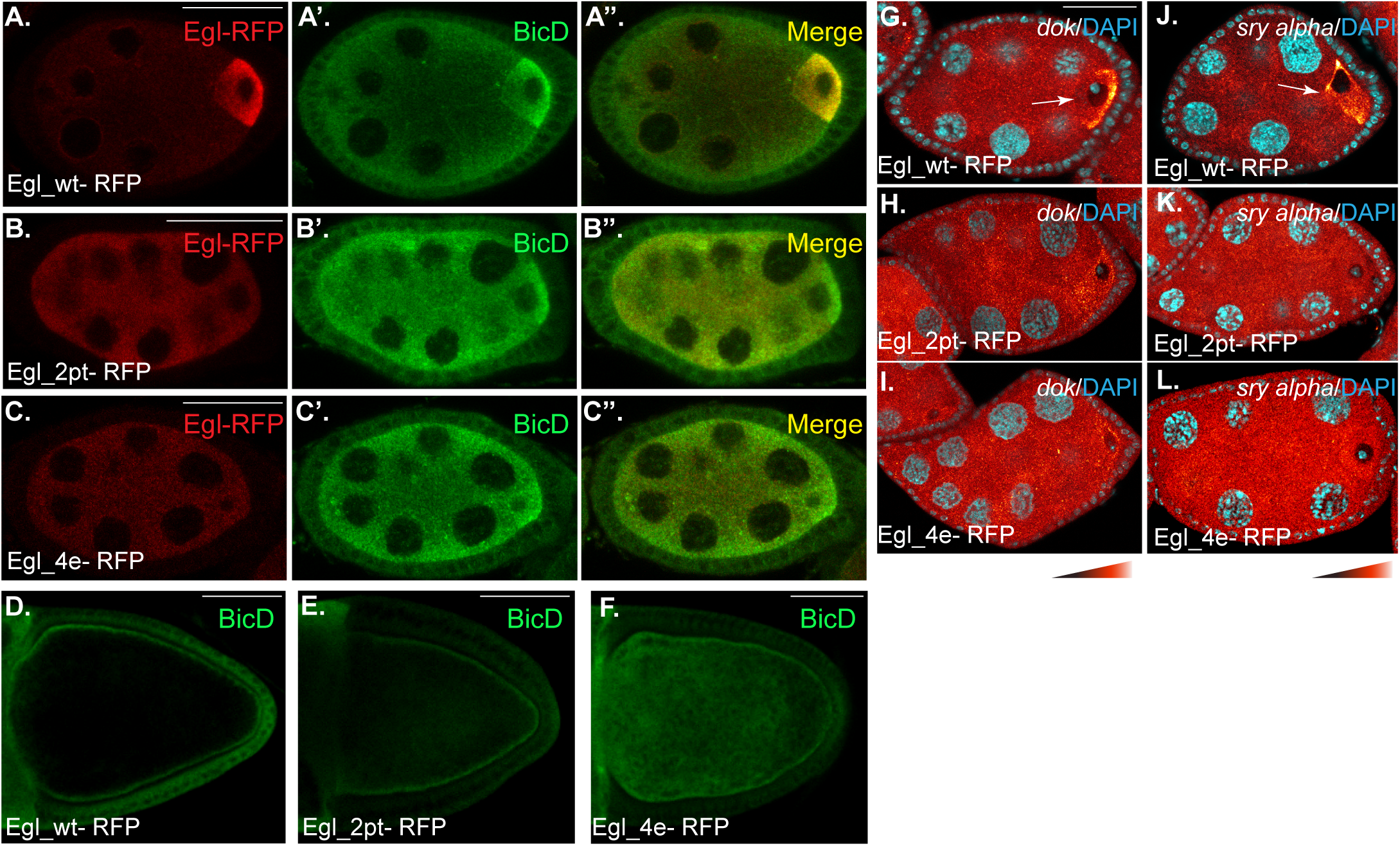
**(A-F)** Ovaries from strains expressing Egl_wt-RFP, Egl_2pt-RFP, or Egl_4e-RFP and depleted of endogenous Egl were fixed and processed for immunofluorescence using an antibody against RFP (red) and BicD (green). Representative stage 5 (A-C) and stage 10 (D-F) egg chambers are shown. **(G-L)** The same strains were fixed and processed for smFISH using probes against *dok* (G-I) or *sry alpha* (J-L). The in situ signal is depicted using a red to white LUT; red pixels represent low-intensity signal and white pixels represent high-intensity signal. The scale bar for D-F represents 50 microns; for the remainder of the images, the scale bar represents 25 microns.

## Bibliography

Barbar, E. (2008) ‘Dynein light chain LC8 is a dimerization hub essential in diverse protein networks’, Biochemistry 47(2): 503–8.

Berleth, T., Burri, M., Thoma, G., Bopp, D., Richstein, S., Frigerio, G., Noll, M. and Nusslein-Volhard, C. (1988) ‘The role of localization of bicoid RNA in organizing the anterior pattern of the Drosophila embryo’, EMBO J 7(6): 1749–56.

Bischof, J., Maeda, R. K., Hediger, M., Karch, F. and Basler, K. (2007) ‘An optimized transgenesis system for Drosophila using germ-line-specific phiC31 integrases’, Proc Natl Acad Sci U S A 104(9): 3312–7.

Bullock, S. L. and Ish-Horowicz, D. (2001) ‘Conserved signals and machinery for RNA transport in Drosophila oogenesis and embryogenesis’, Nature 414(6864): 611–6.

Cajigas, I. J., Tushev, G., Will, T. J., tom Dieck, S., Fuerst, N. and Schuman, E. M. (2012) ‘The local transcriptome in the synaptic neuropil revealed by deep sequencing and high-resolution imaging’, Neuron 74(3): 453–66.

Carpenter, A. T. (1994) ‘Egalitarian and the choice of cell fates in Drosophila melanogaster oogenesis’, Ciba Found Symp 182: 223–46; discussion 246-54.

Cha, B. J., Serbus, L. R., Koppetsch, B. S. and Theurkauf, W. E. (2002) ‘Kinesin I-dependent cortical exclusion restricts pole plasm to the oocyte posterior’, Nat Cell Biol 4(8): 592–8.

Clark, I., Giniger, E., Ruohola-Baker, H., Jan, L. Y. and Jan, Y. N. (1994) ‘Transient posterior localization of a kinesin fusion protein reflects anteroposterior polarity of the Drosophila oocyte’, Curr Biol 4(4): 289–300.

Delanoue, R., Herpers, B., Soetaert, J., Davis, I. and Rabouille, C. (2007) ‘Drosophila Squid/ hnRNP helps Dynein switch from a gurken mRNA transport motor to an ultrastructural static anchor in sponge bodies’, Dev Cell 13(4): 523–38.

Dick, T., Ray, K., Salz, H. K. and Chia, W. (1996) ‘Cytoplasmic dynein (ddlc1) mutations cause morphogenetic defects and apoptotic cell death in Drosophila melanogaster’, Mol Cell Biol 16(5): 1966–77.

Dienstbier, M., Boehl, F., Li, X. and Bullock, S. L. (2009) ‘Egalitarian is a selective RNA-binding protein linking mRNA localization signals to the dynein motor’, Genes Dev 23(13): 1546–58.

Dix, C. I., Soundararajan, H. C., Dzhindzhev, N. S., Begum, F., Suter, B., Ohkura, H., Stephens, E. and Bullock, S. L. (2013) ‘Lissencephaly-1 promotes the recruitment of dynein and dynactin to transported mRNAs’, J Cell Biol 202(3): 479–94.

Ephrussi, A., Dickinson, L. K. and Lehmann, R. (1991) ‘Oskar organizes the germ plasm and directs localization of the posterior determinant nanos’, Cell 66(1): 37–50.

Goldman, C. H. and Gonsalvez, G. B. (2017) ‘The Role of Microtubule Motors in mRNA Localization and Patterning Within the Drosophila Oocyte’, Results Probl Cell Differ 63: 149–168.

Hoogenraad, C. C., Akhmanova, A., Howell, S. A., Dortland, B. R., De Zeeuw, C. I., Willemsen, R., Visser, P., Grosveld, F. and Galjart, N. (2001) ‘Mammalian Golgi-associated Bicaudal-D2 functions in the dynein-dynactin pathway by interacting with these complexes’, EMBO J 20(15): 4041–54.

Kardon, J. R. and Vale, R. D. (2009) ‘Regulators of the cytoplasmic dynein motor’, Nat Rev Mol Cell Biol 10(12): 854–65.

Kim-Ha, J., Smith, J. L. and Macdonald, P. M. (1991) ‘oskar mRNA is localized to the posterior pole of the Drosophila oocyte’, Cell 66(1): 23–35.

Lecuyer, E., Yoshida, H., Parthasarathy, N., Alm, C., Babak, T., Cerovina, T., Hughes, T. R., Tomancak, P. and Krause, H. M. (2007) ‘Global analysis of mRNA localization reveals a prominent role in organizing cellular architecture and function’, Cell 131(1): 174–87.

Li, M., McGrail, M., Serr, M. and Hays, T. S. (1994) ‘Drosophila cytoplasmic dynein, a microtubule motor that is asymmetrically localized in the oocyte’, J Cell Biol 126(6): 1475–94.

Liu, G., Sanghavi, P., Bollinger, K. E., Perry, L., Marshall, B., Roon, P., Tanaka, T., Nakamura, A. and Gonsalvez, G. B. (2015) ‘Efficient Endocytic Uptake and Maturation in Drosophila Oocytes Requires Dynamitin/p50’, Genetics 201(2): 631–49.

Mach, J. M. and Lehmann, R. (1997) ‘An Egalitarian-BicaudalD complex is essential for oocyte specification and axis determination in Drosophila’, Genes Dev 11(4): 423–35.

McClintock, M. A., Dix, C. I., Johnson, C. M., McLaughlin, S. H., Maizels, R. J., Hoang, H. T. and Bullock, S. L. (2018) ‘RNA-directed activation of cytoplasmic dynein-1 in reconstituted transport RNPs’, Elife 7.

McGrail, M. and Hays, T. S. (1997) ‘The microtubule motor cytoplasmic dynein is required for spindle orientation during germline cell divisions and oocyte differentiation in Drosophila’, Development 124(12): 2409–19.

Mili, S., Moissoglu, K. and Macara, I. G. (2008) ‘Genome-wide screen reveals APC-associated RNAs enriched in cell protrusions’, Nature 453(7191): 115–9.

Navarro, C., Puthalakath, H., Adams, J. M., Strasser, A. and Lehmann, R. (2004) ‘Egalitarian binds dynein light chain to establish oocyte polarity and maintain oocyte fate’, Nat Cell Biol 6(5): 427–35.

Neuman-Silberberg, F. S. and Schupbach, T. (1993) ‘The Drosophila dorsoventral patterning gene gurken produces a dorsally localized RNA and encodes a TGF alpha-like protein’, Cell 75(1): 165–74.

Ni, J. Q., Liu, L. P., Binari, R., Hardy, R., Shim, H. S., Cavallaro, A., Booker, M., Pfeiffer, B. D., Markstein, M., Wang, H. et al. (2009) ‘A Drosophila resource of transgenic RNAi lines for neurogenetics’, Genetics 182(4): 1089–100.

Rapali, P., Szenes, A., Radnai, L., Bakos, A., Pal, G. and Nyitray, L. (2011) ‘DYNLL/LC8: a light chain subunit of the dynein motor complex and beyond’, FEBS J 278(17): 2980–96.

Ryder, P. V. and Lerit, D. A. (2018) ‘RNA localization regulates diverse and dynamic cellular processes’, Traffic 19(7): 496–502.

Sanghavi, P., Liu, G., Veeranan-Karmegam, R., Navarro, C. and Gonsalvez, G. B. (2016) ‘Multiple Roles for Egalitarian in Polarization of the Drosophila Egg Chamber’, Genetics.

Sladewski, T. E., Billington, N., Ali, M. Y., Bookwalter, C. S., Lu, H., Krementsova, E. B., Schroer, T. A. and Trybus, K. M. (2018) ‘Recruitment of two dyneins to an mRNA-dependent Bicaudal D transport complex’, Elife 7.

Suter, B. (2018) ‘RNA localization and transport’, Biochim Biophys Acta Gene Regul Mech 1861(10): 938–951.

Theurkauf, W. E., Alberts, B. M., Jan, Y. N. and Jongens, T. A. (1993) ‘A central role for microtubules in the differentiation of Drosophila oocytes’, Development 118(4): 1169–80.

Trovisco, V., Belaya, K., Nashchekin, D., Irion, U., Sirinakis, G., Butler, R., Lee, J. J., Gavis, E. R. and St Johnston, D. (2016) ‘bicoid mRNA localises to the Drosophila oocyte anterior by random Dynein-mediated transport and anchoring’, Elife 5.

Vazquez-Pianzola, P., Schaller, B., Colombo, M., Beuchle, D., Neuenschwander, S., Marcil, A., Bruggmann, R. and Suter, B. (2017) ‘The mRNA transportome of the BicD/Egl transport machinery’, RNA Biol 14(1): 73–89.

Weil, T. T., Parton, R., Davis, I. and Gavis, E. R. (2008) ‘Changes in bicoid mRNA anchoring highlight conserved mechanisms during the oocyte-to-embryo transition’, Curr Biol 18(14): 1055–61.

Zhao, T., Graham, O. S., Raposo, A. and St Johnston, D. (2012) ‘Growing microtubules push the oocyte nucleus to polarize the Drosophila dorsal-ventral axis’, Science 336(6084): 999–1003.

